# Nextflow Pipeline for Visium and H&E Data from Patient-Derived Xenograft Samples

**DOI:** 10.1101/2023.07.27.550727

**Authors:** Sergii Domanskyi, Anuj Srivastava, Jessica Kaster, Haiyin Li, Meenhard Herlyn, Jill C. Rubinstein, Jeffrey H. Chuang

## Abstract

**Highlights:** We have developed an automated data processing pipeline to quantify mouse and human data from patient-derived xenograft samples assayed by Visium spatial transcriptomics with matched hematoxylin and eosin (H&E) stained image. We enable deconvolution of reads with Xenome, quantification of spatial gene expression from host and graft species with Space Ranger, extraction of B-allele frequencies, and splicing quantification with Velocyto. In the H&E image processing sub-workflow, we generate morphometric and deep learning-derived feature quantifications complementary to the Visium spots, enabling multi-modal H&E/expression comparisons. We have wrapped the pipeline into Nextflow DSL2 in a scalable, portable, and easy-to-use framework.

**Summary:** We designed a Nextflow DSL2-based pipeline, Spatial Transcriptomics Quantification (STQ), for simultaneous processing of 10x Genomics Visium spatial transcriptomics data and a matched hematoxylin and eosin (H&E)-stained whole slide image (WSI), optimized for Patient-Derived Xenograft (PDX) cancer specimens. Our pipeline enables the classification of sequenced transcripts for deconvolving the mouse and human species and mapping the transcripts to reference transcriptomes. We align the H&E WSI with the spatial layout of the Visium slide and generate imaging and quantitative morphology features for each Visium spot. The pipeline design enables multiple analysis workflows, including single or dual reference genomes input and stand-alone image analysis. We showed the utility of our pipeline on a dataset from Visium profiling of four melanoma PDX samples. The clustering of Visium spots and clustering of imaging features of H&E data reveal similar patterns arising from the two data modalities.

## Introduction

The Patient-Derived Xenograft (PDX) cancer model^1^ is an in vivo system where a patient tumor sample is engrafted, grown, and passaged in an immunocompromised mouse,^2^ such as NSG or NOD/SCID models. PDX models have been shown to maintain the characteristics of the primary patient tumor and recapitulate patient therapy response.^3–7^ Therefore, PDXs are a powerful tool for preclinical therapeutic testing and *in vivo* study of the biological processes that occur in cancer, and are also valuable in the development of new therapies.^8^ Molecular characterization of PDX models is vital for identifying oncogenic signatures to guide model selection for therapeutic studies. However, characterization of PDX models is a complex task that requires community effort to establish repositories and develop standardized tools for analysis that take into account the co-occurrence of mouse and human cells. The characterization of samples from PDX models has routinely been performed by Next Generation Sequencing (NGS) assays, including whole genome sequencing (WGS), whole exome sequencing (WES), transcriptome sequencing (RNA-seq), and single-cell resolved transcriptome (sc/snRNA-seq). For example, the PDXNet portal^9^ was developed as a resource for coordinating and processing NGS data from PDX models of more than 33 cancer types. Recent technology advances have made spatial approaches such as spatial transcriptomics (ST) and automated analysis of histology images increasingly important in the cancer field, and their analysis in PDXs requires a similar development of tools and standards.

Spatially resolved transcriptome sequencing^10^ enables new investigations of cancer and its microenvironment that can be studied in PDXs. Several types of assays for spatial transcriptomics primarily differ by spatial resolution. To obtain ST from macroscopic regions (or from small regions of interest) guided by the expression of markers of interest DSP GeoMx^11^ can be used. Other ST methods can provide highly multiplexed transcriptome measurements at cellular and sub-cellular resolution (MERFISH^12,13^, CosMx). Other ST assays, e.g., Slide-seq and 10x Visium^14^, employ a grid-based approach with spatial resolution typically between 10 and 100 *μm*. Specifically, the Visium Spatial Gene Expression (Visium) slide is developed by 10x Genomics and has 4992 barcoded spots of 55 μm diameter distributed over a hexagonal lattice.

The Visium ST protocol also allows H&E staining of the same tissue section used to prepare the RNA sequencing libraries, enabling comparative histopathology and expression analysis for PDXs. The H&E-stained slide is scanned for a full-resolution digital Whole Slide Image (WSI). The typical resolution of a WSI is 0.25 *μm* per image pixel. WSI of the Visium slide is inspected and analyzed by a pathologist. The WSI can determine tissue slide quality, detect artifacts, and annotate morphological structures. Manual detailed assessment of the WSI image is a laborious process, and machine learning tools can help to quantify morphological features on WSI^15,16^.

Before determining the nuclear morphometric features of the WSI in a PDX, it is necessary to identify individual cell nuclei via segmentation methods that are effective on both mouse stromal and human cancer cells. The segmentation yields a set of nuclear boundaries, where each boundary can be further characterized to determine enclosed area, perimeter length, orientation, eccentricity, and other quantities. The individual nuclei can be assigned to ST spots based across the image with grid alignment. Since each ST spot may contain multiple nuclei, it is possible to calculate a nuclear morphometric distribution for each ST spot. In addition, the full-resolution image of each spot can be used to generate an additional layer of information made of imaging features that characterize texture, color gradients, and other aspects of each spot’s image.

Automating and streamlining the processing of raw Visium data from NGS sequencing and imaging of PDX-derived histology slides is an important problem for which new tools are necessary. The sequenced libraries of Visium slides from PDX-derived samples contain human and mouse transcripts. The transcripts must be classified as deriving from the host mouse or graft human sequencing reads or discarded from the analysis. To integrate the transcriptomic features with imaging and morphometric features derived from full-resolution image tiles, the H&E WSI must also be aligned and compared with the Visium ST grid. To overcome the challenges above, we developed the Spatial Transcriptomics Quantification (STQ) pipeline based on Nextflow DSL2 language. We designed the pipeline for PDX Visium ST data and H&E stained WSI processing to enable diverse multimodal analysis.

## Design

### Spatial transcriptomics deconvolution

Processing Visium ST raw data derived from PDX samples requires regimented computational infrastructure and tool specification. An engineering challenge is that the standard workflow in the 10x Genomics software tool Space Ranger does not account for species deconvolution of the sequenced PDX libraries. Therefore, the only existing option to process PDX data with Space Ranger is to use a joint human-mouse reference genome and map each sequencing read to both mouse and human reference genomes simultaneously. Our pipeline improves on the existing process by adding an option to run sequencing reads classification (i.e., species deconvolution or host-graft deconvolution) with Xenome, a human/mouse deconvolution method commonly used in PDX sequence analysis, before processing with Space Ranger.

### Data processing for splicing and genomic inference

The primary output of Space Ranger consists of two components: (1) the gene by spot UMI count matrix and (2) image and grid alignment. Additional outputs are the QC metrics of alignment, detected tissue regions, and Binary Alignment Map files (BAM). In STQ, we add engineering functionality to generate data from the BAM files necessary for important downstream analysis tasks, including RNA splicing quantification, lineage tracing, and copy number variation inference.

### H&E image processing

Analysis of the full-resolution H&E images requires a complex set of tools that align the image to the Visium ST grid and extract imaging and nuclear morphometric features from each spot’s image. The feature extraction and image segmentation are done with cutting-edge machine learning methods such as Convolutional Neural Networks (CNN) and other deep learning architectures.

### STQ analysis workflows

We have developed three analysis workflows for data processing with the STQ pipeline. These workflows correspond to three ways the Visium PDX data can be analyzed. The first workflow is designed to analyze PDX samples Visium ST data (**Figure 1**, green lines) by explicitly mapping of mouse reads to the mouse reference transcriptome and human reads to the human reference transcriptome. The analysis starts with Xenome reads classification. The Xenome indices necessary for classification are built by the pipeline automatically in a one-time computation step. For each deconvolved species, the FASTQ reads files are sorted and used as input to the Space Ranger processing step. Space Ranger genome separate mapping of each species’ transcripts and image grid alignment are followed by gene splicing quantification and extraction of bulk-level SNVs. The grid alignment is then used to assign a full-resolution H&E-stained slide image for each ST spot. The imaging and nuclear morphometric features are extracted for each spot’s image, generating the output feature by spot matrix.

**Figure 1.**
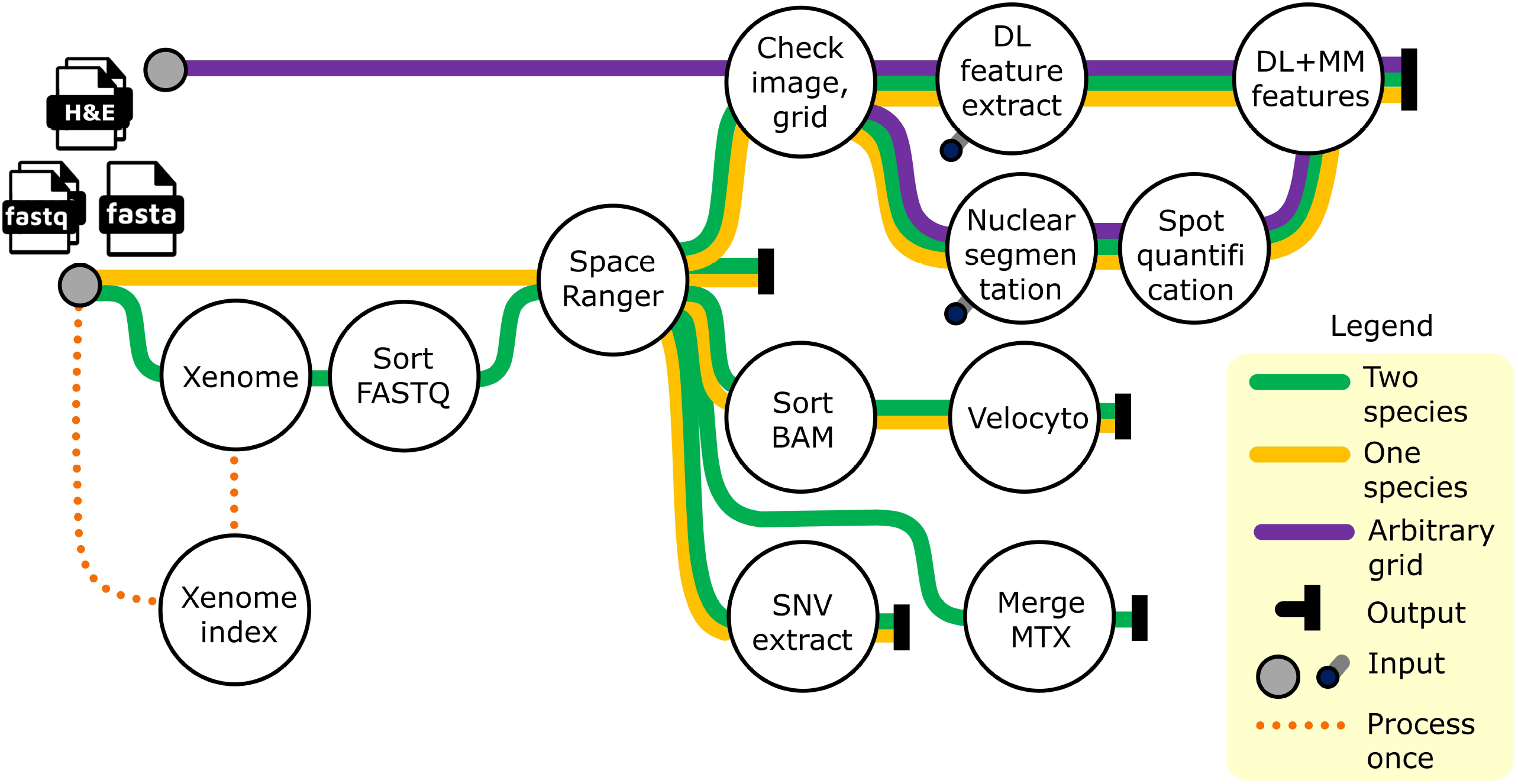
Workflows of analysis. The workflows provide three ways the Visium PDX data, i.e., FASTQ files and an H&E-stained image, can be analyzed with the STQ pipeline. The first workflow (green lines) uses two genome references and allows species classification of sequencing reads supplied as FASTQ files. In this analysis workflow, separate mapping of each species reads to the species’ reference transcriptome with Space Ranger is carried out. The second workflow (yellow lines) uses a single reference transcriptome input and starts the analysis by processing the reads with Space Ranger without species reads deconvolution. The third workflow (purple lines) is a stand-alone full-resolution image processing that disregards the Visium spot grid. Before using the green analysis workflow, Xenome indices must be prepared in a one-time process indicated by an orange dotted line. Depending on the analysis workflow and setup of pipeline options, not all output components may be generated. Pipeline inputs are shown as black grey-filled circles. Additional inputs shown as black circles connected to processes with grey lines are required by the deep learning (DL) and nuclear morphometric (MM) features extraction tools. H&E – hematoxylin and eosin, FASTQ – sequencing reads with quality scores file format, FASTA – reference sequencing reads file format, BAM – binary alignment map file format, SNV – single nucleotide variant, MTX – matrix market coordinate format.

The second analysis workflow (**Figure 1**, yellow lines) is used with a single genome reference and, therefore, for *simultaneous* mapping of sequencing reads from PDX samples to a joint human+mouse reference. The imaging features extraction workflow follows the same steps as in the first analysis workflow. The second analysis workflow can be used for any Visium ST dataset, including cases where the sample contains reads from a single species, e.g., human. The user can specify the desired reference genome for read mapping.

The third analysis workflow, **Figure 1**, purple lines, is designed to generate an arbitrary spot grid, align the full-resolution H&E image to the grid, and extract imaging and nuclear morphometric features.

Each analysis workflow is implemented as a Nextflow DSL2 workflow. Each workflow consists of one or more sub-workflows. **Figure 2** shows a schematic of the sequencing sub-workflow of the two-reference analysis workflow. Reads classified by Xenome as “both,” “ambiguous,” or “neither” are discarded. Human and mouse reads are processed further in the Space Ranger step. Genome-sorted BAM files, produced by Space Ranger, are cell barcode-sorted and processed with Velocyto^17^ to estimate spliced, un-spliced and ambiguous UMI counts for each gene and every ST spot. Bulk-level SNV estimation is generated for each BAM file, proving B-allele frequency (BAF) shifts. Details of BAF data generation are illustrated in **Figure 3**. BAF data can be used with CaSpER for robust inference of CNV from the transcriptomics data. The CaSpER CNV inference is made on integrated data samples, and thus CaSpER CNV inference cannot be included in the STQ pipeline.

**Figure 2.**
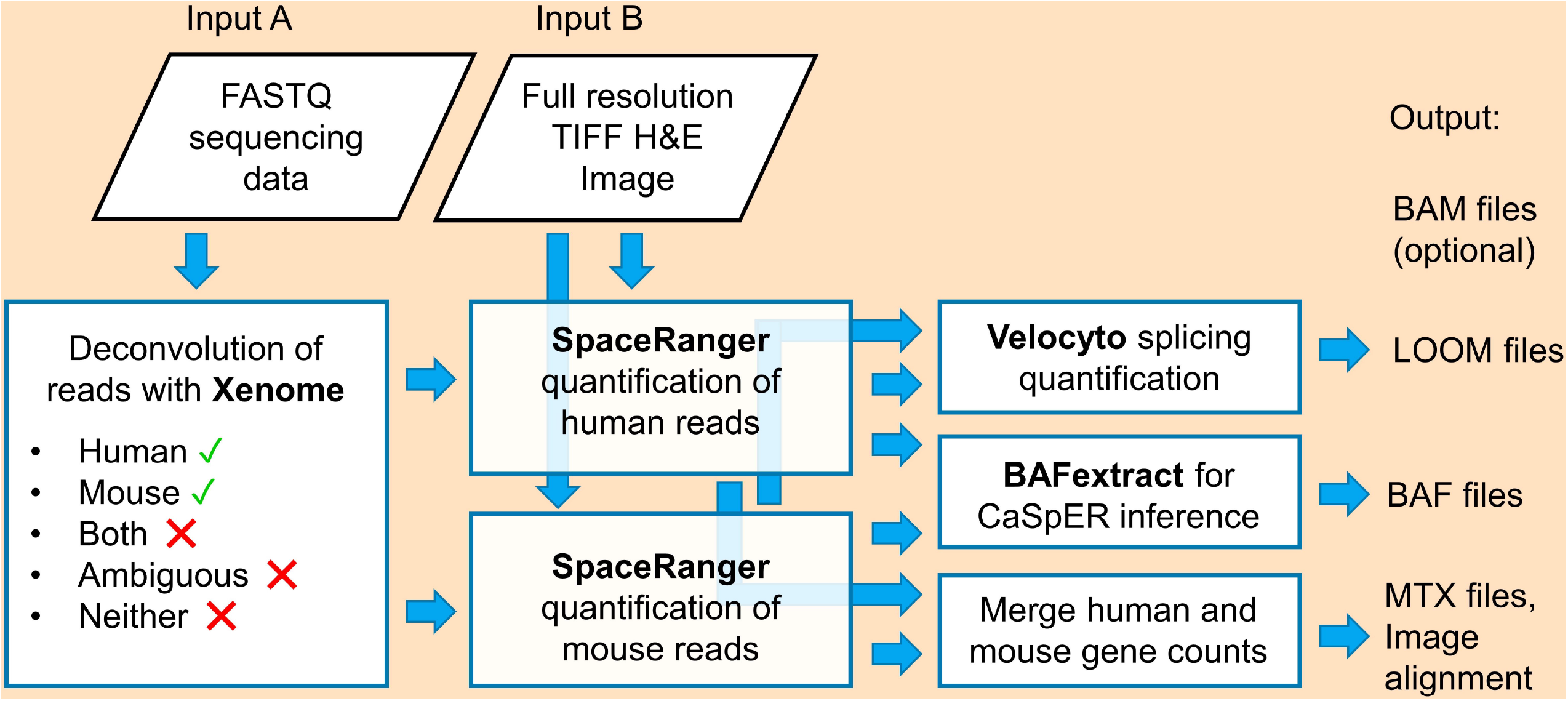
Sub-workflow for Visium sequencing data quantification. The sub-workflow takes input data (input A and input B), mouse and human reference transcriptomes, and Xenome reference indices. The analysis starts with reads deconvolution, genome mapping, full-resolution image, and grid alignment for the two-reference analysis workflow. Splicing quantification with Velocyto, extraction of B-allele frequencies (BAF), and merging of the gene count matrices are performed on Space Ranger outputs. FASTQ – sequencing reads and quality scores file format, TIFF – tag image file format, H&E – hematoxylin and eosin, BAM – binary alignment map file format, LOOM – large omics matrix file format, MTX – matrix market coordinate format.

**Figure 3.**
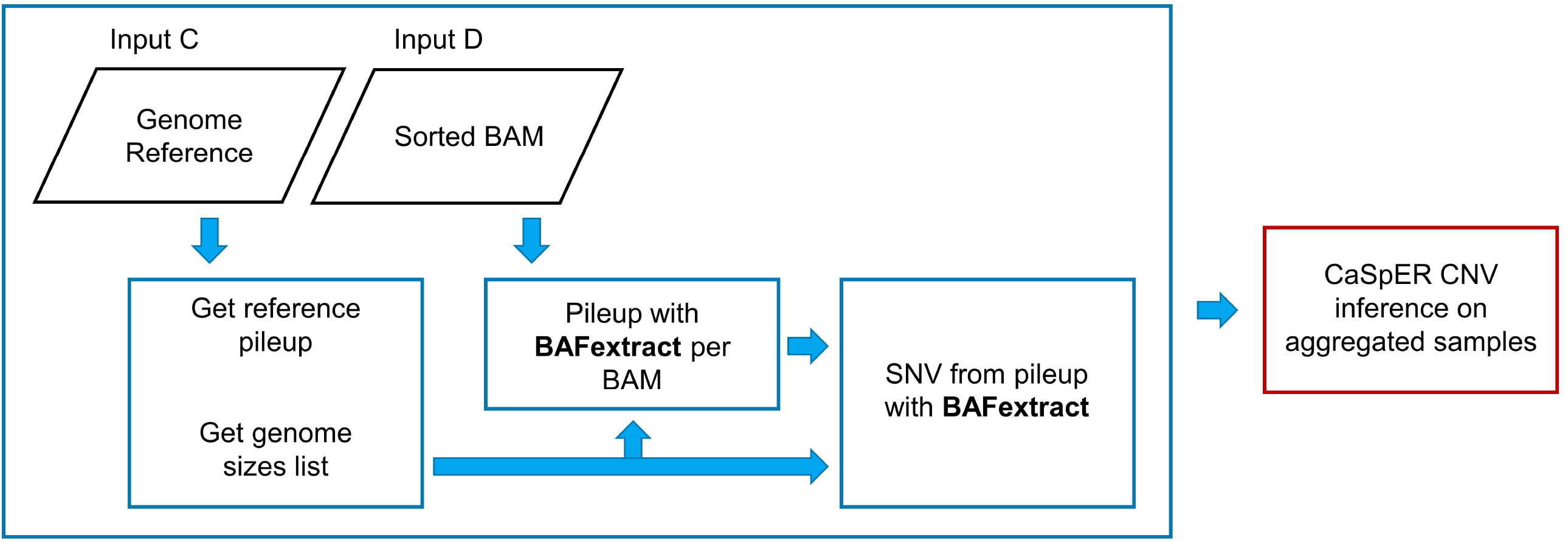
Schematic for BAF (B-allele frequencies) extraction. Detailed view on a BAFextract section of the sub-workflow for Visium sequencing data quantification, where the input consists of parts C and D, the reference genome, and the genome-sorted BAM files produced by Space Ranger. The reference genome is preprocessed, and the tool BAFextract is then used to generate a BAF file for each input BAM file. The red box on the right side depicts the process not included in STQ to illustrate how the BAF data can be used in downstream analysis.

The imaging and nuclear morphometric feature extraction sub-workflow that is a part of all three analysis workflows is illustrated in **Figure 4**. Briefly, a region of interest (ROI) is cut out of the WSI, color and /or stain-normalized, and imaging features are extracted for each spot image (a square tile) on a user-defined or Space Ranger-generated grid. The nuclear segmentation and, optionally, HoVer-Net^18^ classification is performed on the WSI ROI. HoVer-Net is a CNN that uses information encoded within the vertical and horizontal distances of image pixels of nuclei to their centers of mass to do simultaneous segmentation and classification of nuclei from H&E-stained histology images.^18^ The STQ HoVer-Net process uses pre-trained^19^ model weights derived from a large number of histology images to enable accurate segmentation and classification of nuclei. The default nuclear segmentation method of STQ is StarDist^20,21^ which is less computationally-intensive analysis of than HoVer-Net. StarDist is a U-Net-based deep neural network that localizes nuclei via star-convex polygons. In short, StarDist produces an overcomplete set of candidate polygon boundaries for each nucleus and then determines the optimal set of nuclear boundaries. The imaging sub-workflow is completed by generating the averages and standard deviations of nuclear morphometric features for each grid spot.

**Figure 4.**
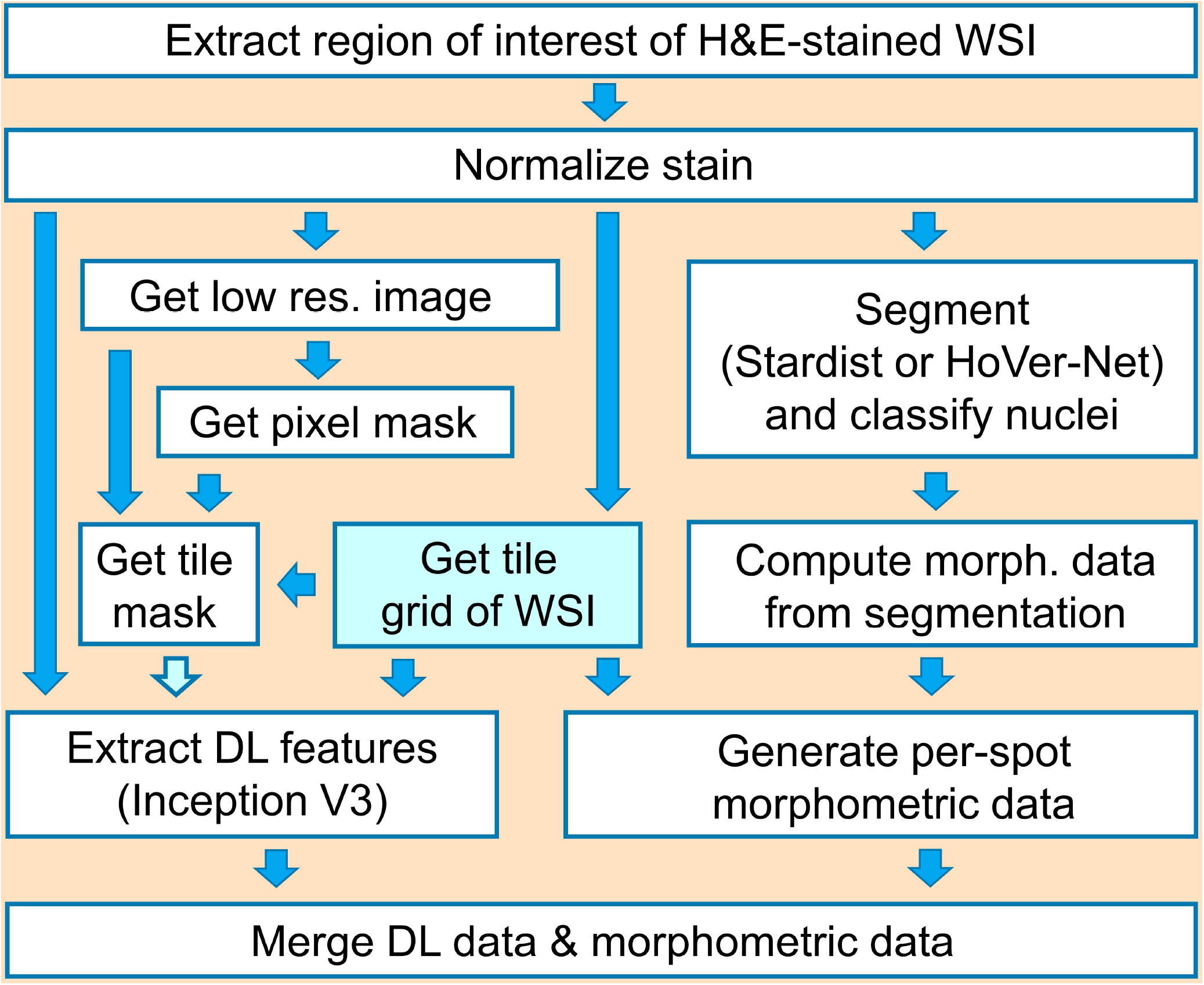
Schematic of the imaging sub-workflow. Full-resolution H&E analysis with deep learning (DL). The first step of the sub-workflow is to extract a region of interest, if specified, and convert the image to a decompressed TIFF image. The major sub-workflow components include image stain normalization, grid preparation, extraction of DL, and nuclear morphometric features. The light blue highlighted process is omitted when using the existing grid generated with SpaceRanger. The extracted DL and morphometric features for each tile is the main output of the imaging sub-workflow. H&E – hematoxylin and eosin, WSI – whole slide image, TIFF – tag image file format, res. – resolution, morph. - morphology.

## Results

To illustrate these workflows, we have applied them to four technical replicates. These are samples from a single Visium slide, where each PDX tissue section originated from the same flash frozen OCT-embedded PDX-derived tumor block (See Methods for experimental details used to prepare the slide). The tumor block was generated from a PDX, model WM4237-1 of BRAF V600E mutant melanoma. We use the four tissue sections derived from the block to develop and test the STQ pipeline. The STQ pipeline generates pre-processed Visium and H&E-derived data for mechanistic and exploratory downstream analysis.

### Clustering RNA-based and Imaging-based features

The transcriptomics output of the STQ pipeline is processed to filter genes by spot count matrices for each sample, normalize UMI counts, and scale library sizes for each sample. We integrate the samples by finding the union of sample-specific highly variable genes, performing PCA, and performing batch correction of principal components. We construct a k-nearest neighbors’ graph and determine clusters of spots (see methods for parameter values). The spatial layout of the transcriptomics data, where each Visium spot is color-coded to the cluster identifier, is shown in **Figure 5A**.

**Figure 5.**
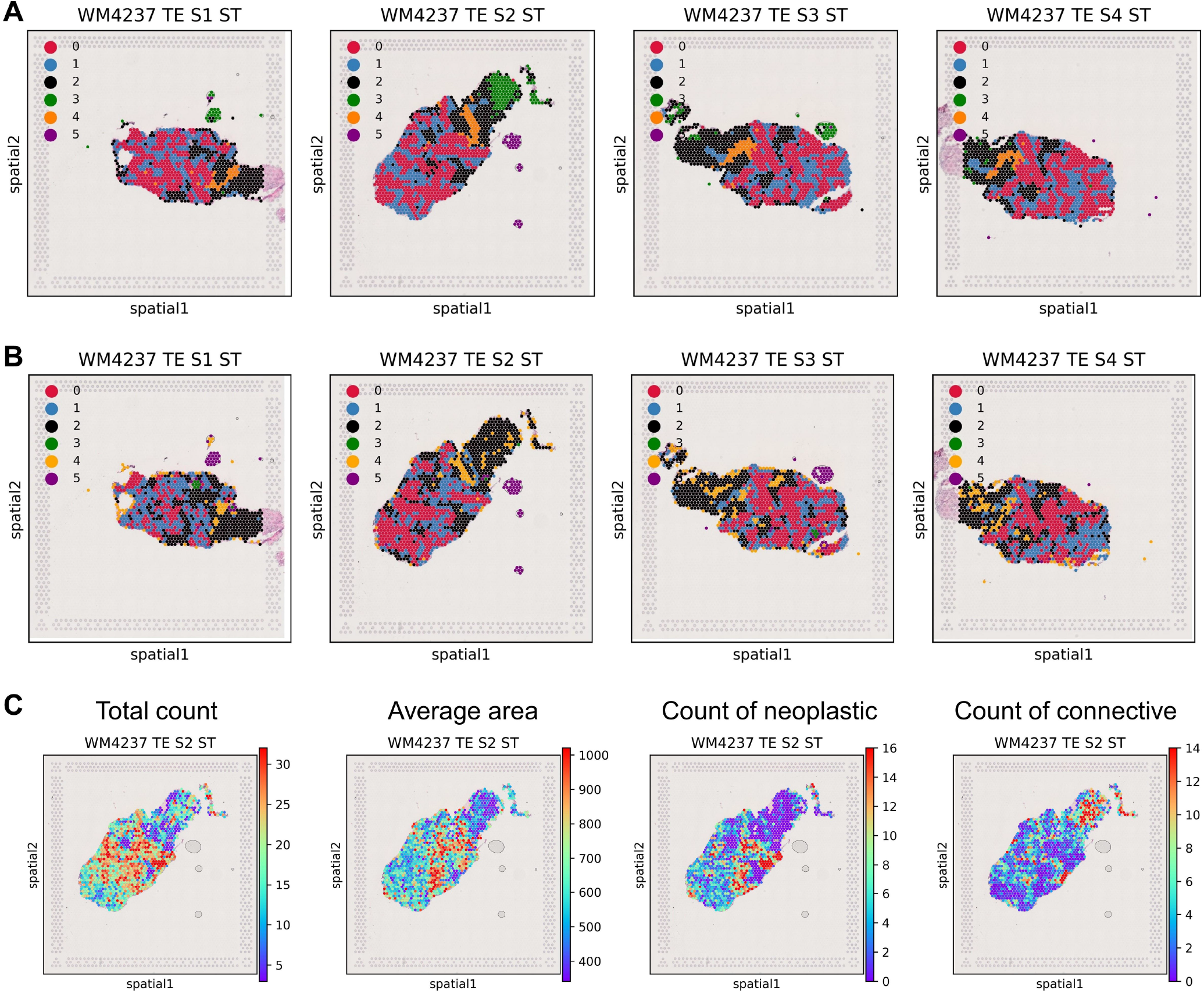
Downstream analysis of the STQ pipeline transcriptomic and imaging output from a Visium dataset. Each sample in the dataset is a spatially barcoded whole transcriptome sequencing and a matched H&E-stained PDX tumor tissue section image on a Visium slide capture area. (A) Visualization of clusters derived from RNA quantification followed by quality control, dimensionality reduction, and clustering of the Visium spots, and (B) H&E-derived clusters of spots based on deep learning (DL) imaging features. (C) An example of the results of nuclear segmentation of the H&E image of sample S2 aggregated on Visium spots to derive the total per-spot nuclei count and the average nucleus area in each Visium spot shown in the two left panels. The two right panels show the result of HoVer-Net nuclei classification, aggregated to the spot level to derive nuclei counts per Visium spot for neoplastic and connective cells. H&E – hematoxylin and eosin.

We also analyze 2,048 imaging features extracted from each spot’s full-resolution H&E image. After determining highly variable imaging features, we perform PCA and correct the principal components to reduce batch effects across the four samples. As in the transcriptomics analysis, we build a k-nearest neighbors’ graph and determine preliminary clusters of spots. To facilitate visual comparison, after the preliminary clustering, we re-map the imaging cluster identities to approximately align their colors with the colors of the RNA-based clusters; specifically, we swap the imaging cluster identities 0,1,2,3,4,5 with 2,0,1,4,5,3. The spot cluster identities derived from the imaging features can be projected onto the tissue image, **Figure 5B**.

### Nuclear morphometric features

The imaging sub-workflow of the STQ pipeline enables nuclear segmentation and classification. Each nucleus boundary and assigned label is saved in the pipeline output directory. The average nuclear morphometric quantities are derived for each spot, including total nuclei count, average nucleus area, average nucleus perimeter length, average nucleus major axis orientation, and average nucleus eccentricity. **Figure 5C** shows the total count and average nucleus area for each spot of sample S2. Standard deviations of the nuclear morphometric quantities are generated as well. Since nuclei are classified into several classes (e.g., neoplastic, connective, inflammatory, and necrotic), the morphometric features and their standard deviations are also calculated for each nucleus class. See **Figure 5C** for examples of neoplastic and connective nuclei counts in sample S2.

### Alignment to separate reference transcriptomes of mouse and human accounts for ambiguous reads

The “One species” analysis workflow uses a combined human+mouse reference transcriptome, which can be downloaded from the 10x Genomics portal.^22^ Each gene name in the reference is prefixed with the species identifier. The Space Ranger step of the STQ pipeline maps each sequenced read to the combined reference transcriptome.

The “Two species” analysis workflow is designed to take two reference transcriptomes, e.g., mouse and human reference, and perform read classification with Xenome^23^ before the Space Ranger read alignment step. Reads mapped to both species are classified as ambiguous, and unmapped reads are filtered out. For reads classified as host, the Space Ranger reads alignment is done using the mouse reference transcriptome. For reads classified as graft, alignment is done using the human reference transcriptome. This “Two species” analysis workflow for analysis of the PDX-derived sequencing reads explicitly accounts for ambiguous reads homologous between host and graft species in the PDX model.

## Discussion

This application of the STQ pipeline to melanoma PDX 10x Visium data of untreated flash frozen sections shows the versatility of analysis for PDX spatial gene expression and H&E-stained WSIs. Depending on the selection of downstream analysis tasks, the user can configure the STQ pipeline to generate desired outputs. The output files have a standardized format to facilitate use with other research computational tools and software packages.

### Limitations

To execute the STQ pipeline efficiently and avoid potential data processing bottlenecks when analyzing multiple samples, the user should have a High-Performance Computing (HPC) infrastructure available. The pipeline has been optimized with the HPC workload scheduler Simple Linux Utility for Resource Management (SLURM). However, a custom pipeline configuration can be prepared for systems using a scheduler other than SLURM. For example, for running the STQ pipeline on the Google Cloud Platform using a beta engine of Google Life Sciences, the user can create a Google storage bucket and set up a Google Cloud Shell to install Nextflow, Singularity, and the STQ pipeline. After updating the pipeline configuration to include a new profile with the Google Life Sciences compute engine as the process executor, the user can process the Visium and H&E data from their Google Storage bucket. Alternatively, instead of using Google Life Sciences compute engine, the user can deploy a SLURM cluster compute engine based on the Google Platform and add a Google SLURM profile to the STQ pipeline configuration. If no grid scheduler is available, then the STQ pipelines can be configured and executed in a local mode, provided that the executor machine has sufficient CPU, memory, and storage to complete each pipeline step.

The deep learning tools in the imaging sub-workflow of the STQ pipeline have been implemented for use with CPUs. Even though many individual tools would run faster in GPU-enabled environments, the parallel execution of the pipeline processes in CPU-only environments allows excellent scalability.

Nearly each STQ pipeline step requires using a specialized tool provided in a software container. We used singularity containers for all tools and built the pipeline configuration on singularity. Some research communities, such as Nf-core,^24^ may require bioconda, docker, or other types of software containers.

## Conclusions

We have presented a Nextflow DSL2 Spatial Transcriptomics Quantification pipeline for processing 10x Genomics Visium Spatial Gene Expression Slide data: spatially barcoded sequencing reads and a full-resolution H&E stained WSI of the capture area. The pipeline is designed for use with ST data generated from PDX-derived samples, where the sequencing reads must first be classified into the mouse and human portions and then aligned to the respective mouse and human reference transcriptomes. Alternatively, the STQ pipeline can process all reads from the PDX-derived ST data and map them to a common mouse-human reference transcriptome.

The STQ pipeline can be used with any single species sample generated from the Visium slide with the “One reference” analysis workflow. For example, a patient-derived specimen Visium dataset can be processed with a human reference transcriptome, or a mouse specimen can be processed with a mouse reference transcriptome in this analysis workflow.

If it is necessary to perform an imaging analysis workflow with grid dimensions different from the Visium grid, e.g., a square grid of tightly laid out tiles, the new grid will be generated in the imaging sub-workflow. The pipeline can be used in the “Arbitrary grid” analysis workflow to skip the sequencing sub-workflows.

All three analysis workflows are implemented with scalable and robust processes. The pipeline is highly configurable and can be set up to run in any HPC environment where Singularity and Nextflow software is installed. STQ thus provides a powerful set of engineering tools for analysis of spatial transcriptomics data in PDXs, which are crucial systems for cancer research and clinical trial evaluation.

## Star Methods

### Nextflow DSL2 and Singularity software containers

Nextflow workflow language developed by Seqera Labs is a domain-specific language (DSL) for creating data processing pipelines. Version 2 of Nextflow, Nextflow DSL2, extends the capabilities of Nextflow and simplifies writing complex pipelines with the utility of modules and sub-workflows. We built sub-workflows for sequencing reads processing with one reference, sequencing reads processing with two references, full-resolution image analysis, and Xenome index preparation. The sub-workflows are combined into Nextflow workflows, which we define as analysis workflows, **Figure 1**. The sub-workflows consist of Nexflow DSL2 processes with defined inputs, outputs, and operations. Most of our pipeline processes require a dedicated Singularity software container. The containers extend the toolbox of the host operating system where the pipeline is invoked to include such tools as “fastq-tools”, “xenome”, “spaceranger”, etc., and custom Python environments with a fixed list of packages installed along with required system library dependencies.

### Xenome reads classification

The tool “xenome classify”^23^ requires indices generated by the tool “xenome index” as input. The STQ pipeline attempts to use the indices provided in the pipeline configuration path; however, if the indices are unavailable, the STQ pipeline executes “Xenome index” sub-workflow to prepare indices. The indices used in our analysis were built with xenome_kmer_size 35, custom-built xenome_reference_host from NOD assembly, and xenome_reference_graft from T2T-CHM13v2.0 assembly of GenBank version GCA_009914755.4.^25^

Xenome is designed to classify xenograft-derived RNA-seq reads to deconvolve the graft (human) from the host (mouse) reads. Xenome defines classes of reads: definitely human, probably human, definitely mouse, probably mouse, both, ambiguous, neither. In Xenome, classes definitely human and probably human are combined into the human class; classes definitely mouse and probably mouse are combined into the mouse class. We discard reads classified as both, ambiguous, and neither. The statistics of Xenome reads deconvolution are generated in the sample output directory in the file “xenome.summary.txt”.

### Space Ranger

The tool “space ranger count” is developed by 10x Genomics to process one capture area of a Visium Spatial Gene Expression Slide for flash frozen (FF) and formalin fixed paraffin embedded FFPE tissue samples. We use Space Ranger built-in automatic capture area image alignment, where fiducials and the grid are detected from the digital image of a capture area. The image alignment is independent of the reference transcriptome and the sequencing parameters. The image alignment result is generated in the sample output folder under “spatial” and contains a set of files passed along to the downstream steps of the STQ pipeline.

### Velocyto

Velocyto tool “Velocyto run10x” is run for all 4,992 spots in a Visium ST slide using a reference transcriptome and binary alignment map (BAM) file from Space Ranger output. The raw output BAM file is sorted by position. STQ sorts the BAM to generate barcode-sorted BAM data required by Velocyto. The output is a “velocyto.loom” file in the sample output directory, the species sub-directory when used with the “Two references” analysis workflow, and the sub-directory “baf” when used with the “One references” analysis workflow. The *.loom files can be loaded with velocyto.py or scVelo (https://scvelo.readthedocs.io/)^17^ or other compatible toolsets for analysis of RNA velocity.

### Bulk-scale B-allele frequencies

BAFextract is a tool designed by the authors of CaSpER^26^ and is intended to extract B-allele frequencies (BAF) from BAM files to estimate CNV events in the downstream steps with CaSpER. BAFextract also includes DNA scaffolds and mitochondria (MT) DNA if those are present in the species genome reference. The output is a file “extracted.baf” generated for each input species sub-directory of the sample output directory.

### Stain normalization

The STQ pipeline has optional stain or color normalization. The stain normalization is implemented for use with WSI according to Macenko method^27^ and StainTools^28^, where the stain matrix is obtained from a reference image and from each patch of WSI. Then the stain transformation is applied to all patches of the WSI.

The second option in the STQ pipeline is color normalization via StainNet^29^, which applies pixel-to-pixel transformation. The StainNet pre-trained on histopathology dataset was shown to improve AUC ROC compared to Macenko stain normalization in the classification tasks with deep learning methods.

### Image tiling

The default setting is to generate a grid of square tiles with the same geometry as 10x Visium Slide, except that tiles are set to be square instead of the round shape of ST spots. Note that in this default setting, tiles are not covering the image entirely. Users can change grid parameters in the “analysis.config” file. The tiling grid can be specified as “square”, “hex” or “random”. The horizontal center-to-center tile spacing and tile size are the configurable pipeline parameters.

### Imaging feature extraction

Inception^30^ v3 is a convolutional neural network model developed and trained by Google on more than a million images from the ImageNet^31^ database. In our pipeline, the Inception v3 step is designed to take tiling grid information from a whole slide image in the same format as generated by the Space Ranger pipeline for a 10x Visium ST image. The 2,048 imaging features are generated for each image tile defined by the provided tiling grid.

### Nuclear segmentation

We set the default segmentation method to StarDist^20,21^, a fast and scalable WSI method for segmenting star-convex shapes. In application to the H&E-stained images, StarDist detects hematoxylin-stained nuclei and determines their boundary. After segmentation, we compute moments, *μ*, of the nucleus boundary and derive nuclear morphometric features: area (*A*=*μ*_00_), perimeter length (*L*=∑*l*_*ij*_), angle of orientation of the major axis 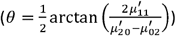, and eccentricity, 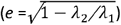, where

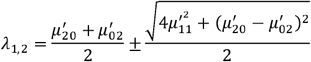

and

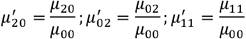

### Nuclear classification

When using the STQ pipeline option of HoVer-Net^18^ nuclear segmentation in the STQ segmentation step, the algorithm predicts a cell-type class for each detected nucleus. The model pre-trained on PanNuke^19^ for HoVer-Net has the following built-in classes: neoplastic, connective, inflammatory, necrotic, not neoplastic, and not labeled. Each assigned label class probability is also stored in the pipeline output.

### Calculation of per-spot nuclear morphometrics

First, each segmented nucleus is assigned to the closest tile on the tile grid. Nuclei outside the tile or Visium ST spot are discarded. The tile assignment boundary can be square or circular and have size determined by the factor parameter in the STQ configuration. We calculate nuclear morphometric features for each nucleus. Then, we calculate the nuclear morphometric features’ average and standard deviation for each tile or spot.

### PDX model generation and sample harvesting

The PDX model WISTAR WM4237-1^32^ is a BRAF V600E mutant metastatic melanoma, subcutaneously implanted solid tumor on NSG mouse #1337, passage 0. The tumor was grown untreated to 300 mm^3^ volume, harvested, and flash frozen, embedded in optimal cutting temperature compound (OCT).

### 10x Genomics Visium Assay generation

cDNA libraries were constructed and sequenced for each sample using the 10x Visium protocol.^33^ The Jackson Laboratory Scientific Services, Single Cell Biology Laboratory generated the assay.

### Full-resolution image acquisition

The 10x Visium ST slide with H&E-stained specimens was digitized with a Hamamatsu C9600-12 NanoZoomer scanner at a resolution of 0.22075 microns per pixel (mpp). The 3-channel WSI was split into four images corresponding to capture areas of the ST slide and converted to TIFF format.

### Gene expression downstream analysis

To explore and inspect the STQ pipeline output for the 4 PDX samples, we apply filtering to keep all expressed genes and ST spots with at least 500 UMI counts and at least 250 genes expressed. We apply the variance stabilizing transformation implemented in the Seurat^34^ package and correct the spot by gene UMI count matrix with the transformation for each of the four samples. We scale each spot library size to 20,000 UMI counts. Using the package scanpy^35^ we determine the top 2,500 highly variable genes (HVG) and perform principal component analysis (PCA) on centered HVG to obtain 50 principal components (PCs). Then we correct the batch effects across the four samples with harmonypy^36^, a Python implementation of the batch effects correction algorithm Harmony^37^. We build a k-nearest neighbors (kNN) graph from the harmony-corrected PCs and cluster spots from all four samples. The color-coded identities of each cluster are projected on the spatial layout of each of the four samples, **Figure 5A**.

### Imaging features downstream analysis

We perform dimensionality reduction and clustering of the ST spots based on their 2,048 imaging features, similar to the gene expression analysis. We determine the top 500 highly variable features (HVF) and perform principal component analysis (PCA) on centered HVF to obtain 50 principal components (PCs). Then we correct the batch effects across the four samples with harmonypy^36,37^. To obtain six clusters, we build a kNN graph from the harmony-corrected PCs and cluster spots from all four samples. The color-coded identities of each cluster are projected on the spatial layout of each of the four samples, **Figure 5B**.

## Data and code availability

The STQ pipeline release v0.1.0 codebase and analysis scripts are hosted at The Jackson Laboratory GitHub: https://github.com/TheJacksonLaboratory/STQ. Raw and processed 10x Visium Spatial Gene Expression data are deposited at Gene Expression Omnibus (GEO) under accession GSE238004, project PRJNA997362. The full-resolution images can be viewed or downloaded at https://images.jax.org/webclient/?show=dataset-4052.

## Acknowledgments

J.H.C. acknowledges support from grants NCI U24-CA224067, R01CA230031 and P30CA034196.

## Author contributions

S.D., A.S., M.H., J.C.R, and J.H.C. conceived the idea for the project; S.D., J.C.R., and J.H.C. worked on the design of the pipeline and downstream analysis of the data. S.D. worked on the development of the pipeline in the nextflow DSL2 framework with support from A.S. All authors provided critical feedback and helped shape the analysis, development, and manuscript. J.K. and H.L. executed the experiment. J.H.C performed the overall supervision of the project.

---

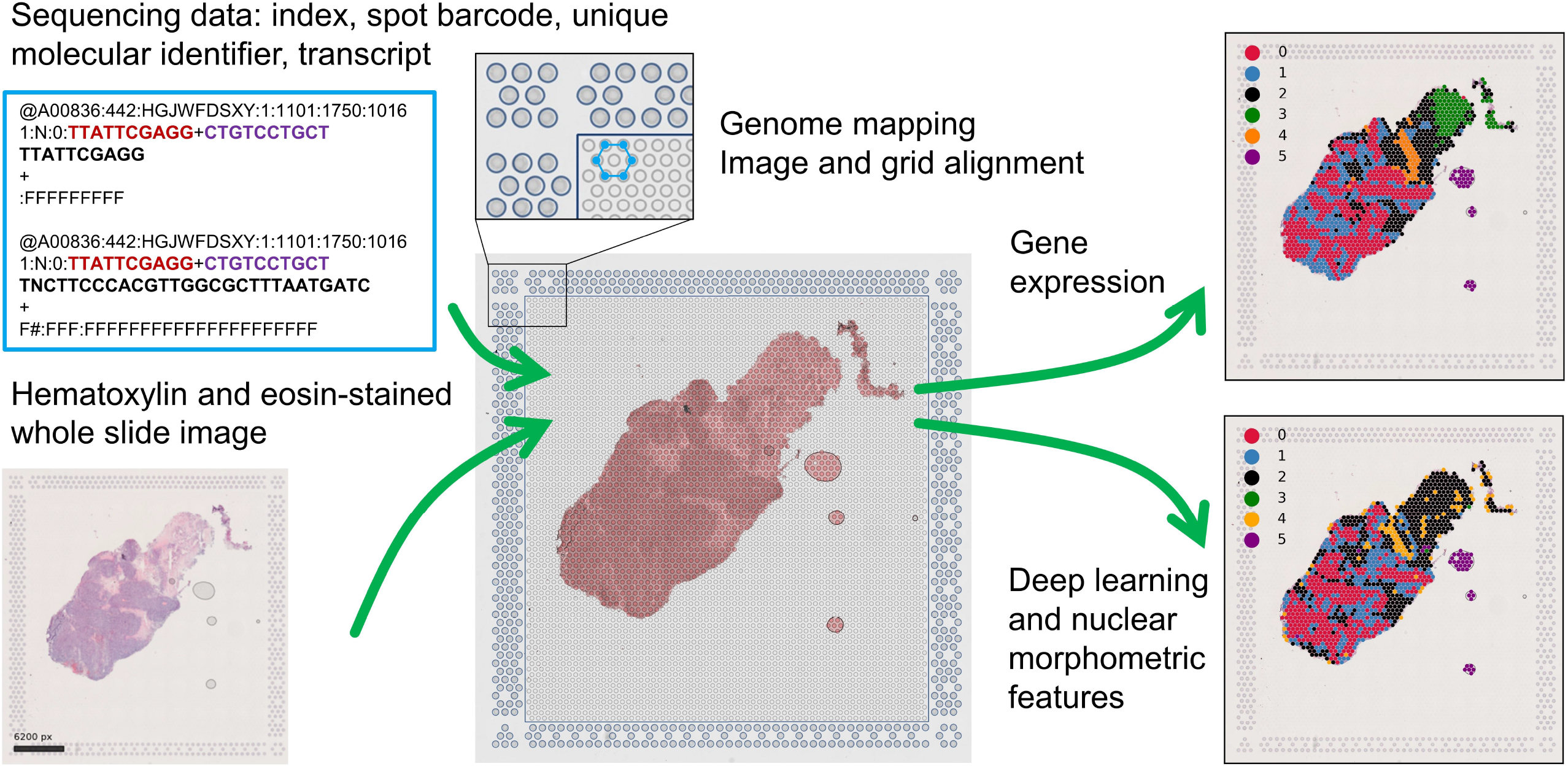

